# Is poor dose selection undermining the translational validity of antidepressant research involving animal models?

**DOI:** 10.1101/2025.10.25.684561

**Authors:** Dasha Anderson, Justyna Hinchcliffe, Megan Jackson, Emma Robinson

## Abstract

**Background:** Behavioural studies in animal models represent a critical component of psychiatric drug development. Positive results in animal studies have identified novel therapeutic targets for major depressive disorder (MDD) but efficacy in humans has largely not been borne out in clinical trials. A possible reason for this failed translation is inappropriate dose selection and the engagement of mechanisms not directly relevant to antidepressant effects in patients.

**Methods:** We first used PubMed to identify preclinical rodent studies in two assays used to assess antidepressants; the conventional forced swim test, (FST) and more recently developed affective bias test, (ABT). Dose ranges were extracted, as well as information about subjects, timing and route of administration, and justification and efficacy of dose(s). Dose ranges were compared against calculated animal equivalent doses.

**Results:** The median FST dose across all antidepressants was 10mg/kg, with median doses for each drug exceeding the relevant animal equivalent dose by 1.5-25x. In contrast, effective doses in the ABT showed closer alignment to those used clinically. In the second study, 232 ketamine and 202 fluoxetine papers involving MDD-related research in rodents were reviewed. The median dose was 10mg/kg for both drugs, exceeding animal equivalent doses by 1.6-3.2x and 3.2-6.5x for ketamine and fluoxetine, respectively.

**Conclusions:** The results indicate pervasive use of antidepressant doses in conventional models of MDD that may not correspond with doses used in clinical practice. We discuss the implications of using doses which exceed therapeutic levels and the potential to engage receptors and underlying mechanisms which are not relevant to clinical effects.

## Introduction

Psychiatric disorders are highly prevalent [1], contribute most to the global burden of disease [2] and carry a high social and economic cost [3]. Recent years have seen little progress in elucidating the pathophysiology of these disorders and developing novel therapies [4]. Behavioural studies in animal models are a core component of psychiatric research, (i) producing animal ‘models’ of diseases to investigate underlying neurobiological mechanisms, and (ii) predicting the efficacy of novel drug candidates for further investigation [5]. However, a critical factor underlying the lack of progress in psychiatric research is that the majority of novel drugs promising in preclinical studies, are unsuccessful in clinical trials [6]. This failure rate is higher for psychiatric drugs than other drug classes [7].

Several reasons for failed translation from preclinical to clinical studies have been suggested, including poor reproducibility [8], small sample size and underpowered studies [9, 10], limitations in study design and data analysis [11], publication bias [12, 13], and the limited validity of animal models [14]. However, a possibly overlooked factor could be dose with the potential for high doses to engage off-target receptors and mechanisms not directly relevant to the human condition.

Our research developed behavioural readouts for rodents adapted from tasks used to investigate neuropsychological impairments in MDD, specifically affective state-induced biases in cognition (referred to as affective biases) [15]. Pharmacological studies performed in rats in both the judgement bias task (JBT) (affective biases and decision-making) and ABT (affective biases associated with reward learning and memory) suggested lower effective doses of conventional and rapid-acting antidepressants (RAADs) than those reported for traditional models of depression [16]. Affective bias modification has been localised to key brain regions associated with emotional behaviour and vulnerability to negative affective biases is associated with early life adversity [17–20]. These findings support the improved translational validity of the ABT but also raise questions about the reason for these differences. Comparing effective doses in the ABT and clinical equivalent doses for rodents, there is closer alignment than observed for the FST. Most antidepressant drugs show some selectivity for the target receptor but, with increasing doses, will interact with other receptors. The dose-response effects at the target receptor are also important as the occupancy levels achieved result in different functional outcomes. If doses used in animal studies exceed the occupancy of the target receptor achieved in patients and/or engage populations of receptors not relevant to their clinical effects, this will limit the translational validity of the arising behavioural effects.

There are species differences in drug absorption, distribution, metabolism, elimination and bioavailability [21], which make identifying clinically relevant animal doses challenging and mean a simple weight adjustment may not be appropriate. A useful method of converting between human and animal doses is to calculate equivalent doses based on weight and body surface area. Current guidelines for dose conversion between animals and humans suggests the following equation by Nair and Jacob [22]:

AED (mg/kg) = Human dose (mg/kg) x Km ratio

where AED is the animal equivalent dose and the Km ratio is a correction factor which normalises the dose based on body surface area (m^2^), known as allometric scaling.

In this paper, we undertook two illustrative systematic reviews of dose ranges used in preclinical antidepressant research: 1) a comparison of dose ranges used in two behavioural assays of depression (FST and ABT) and 2) a comparison of dose ranges used in behavioural studies testing the conventional antidepressant, fluoxetine or rapid-acting antidepressant, ketamine. For each review we include the AED calculated using the equation above, as well as data for receptor affinities and pharmacokinetic data for fluoxetine and ketamine. We also assessed what rationale was provided for the choice of dose or dose range. A more detailed discussion of the rationale for the choice of behavioural tests and drugs included for this review is provided in the supplementary methods.

## Methods and materials

### Systematic review 1 – A comparison of dose ranges used in the Affective Bias Test and Forced Swim Test

#### Search strategy, selection process, inclusion criteria

All published ABT studies of acute effects of antidepressants were included in this systematic review. The range of antidepressants tested using the ABT dictated the antidepressants included in the FST systematic review, thus enabling a direct comparison of dose ranges used in both tasks. FST studies of fluoxetine and ketamine were taken from the studies identified in systematic review 2. Papers were filtered to isolate those where the FST was performed in rats utilising an acute dosing schedule. For the remaining antidepressants, a PubMed search of FST studies was conducted on October 2021 using the following search strategy:

*(rat) AND ((forced swim test) OR (FST)) AND ((agomelatine) OR (citalopram) OR (chlorimipramine) OR (clomipramine) OR (duloxetine) OR (mirtazapine) OR (reboxetine) OR (sertraline) OR (venlafaxine) OR (vortioxetine)).*

In accordance with the PRISMA guidelines [23], an initial screen of titles and abstracts was performed. Full texts of papers that passed this initial screen were then assessed for final inclusion (**Figure 1**). Full text inclusion criteria are listed in **Table 1**.

**Figure 1.**
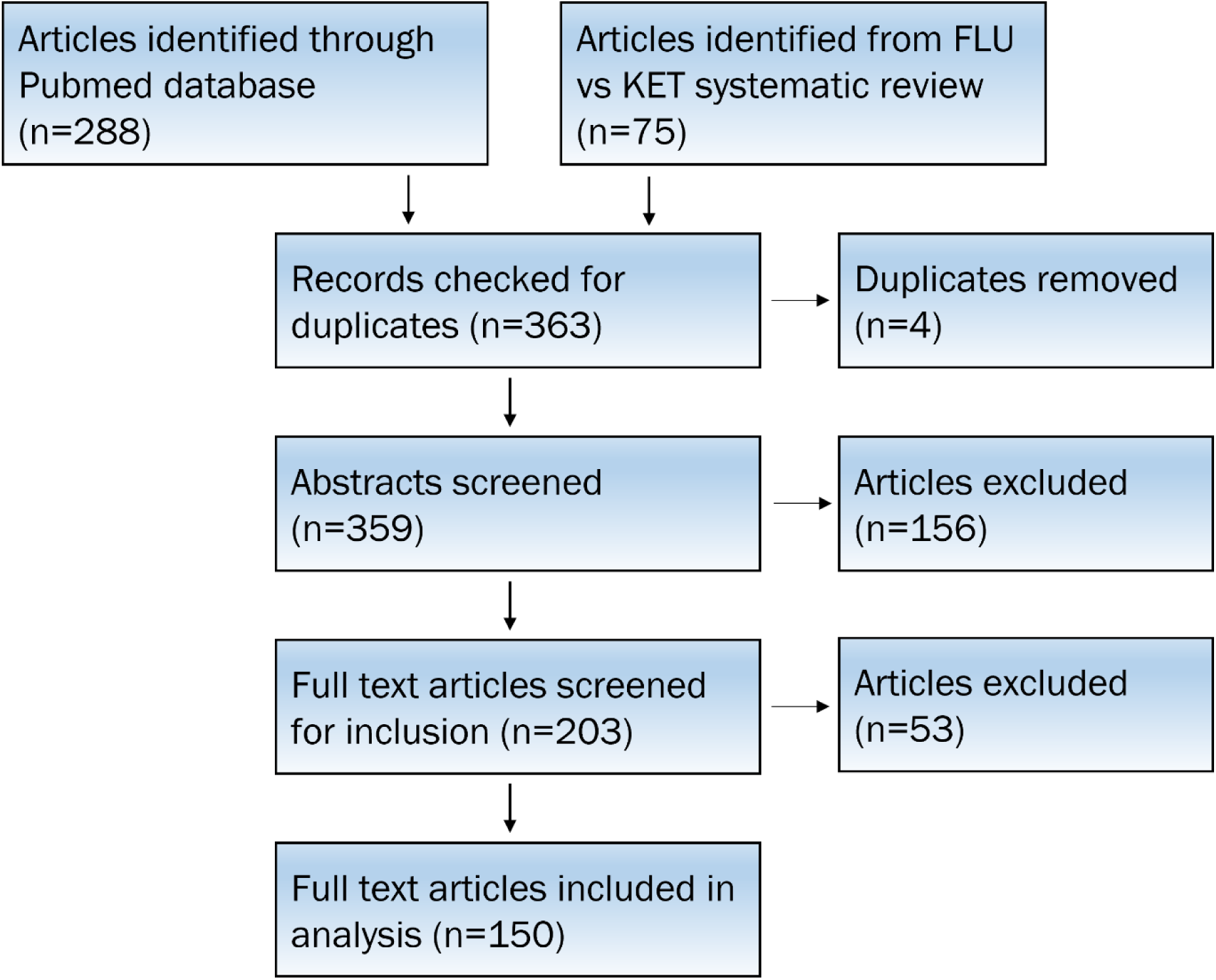
Selection of studies for inclusion in FST systematic review.

**Table 1.** Full text inclusion criteria for FST systematic review.

| Inclusion criteria: |
| --- |
| • Drug administered systemically<br>• Reported route of administration, dose, and behavioural readout<br>• Not a review or meta-analysis<br>• English language paper<br>• Paper directly accessible from PubMed<br>• Conducted in rats<br>• Examined FST<br>• Examined antidepressant previously studied using the ABT (agomelatine, citalopram, clomipramine, duloxetine, fluoxetine ketamine, mirtazapine, reboxetine, sertraline, venlafaxine, and vortioxetine)<br>• Examined acute dose administration |

#### Data extraction and summary

The main outcome measure was the range of doses used, which was compared against the mouse and rat AEDs calculated for each. The AED for ketamine was calculated using the dose from the original clinical trial demonstrating its clinical efficacy [24] and for other antidepressants the clinical dose was taken from the British National Formulary database. Several secondary outcome measures were also extracted, including the species, sex and strain of the animals which were used to convert doses into mg/kg where necessary (**Table 2**). Dose efficacy in FST studies was defined as a significant reduction in immobility, climbing and/or swimming time and dose efficacy in ABT studies was defined as a significant positive choice bias toward the drug-paired substrate. Where multiple administration methods or dosing regimens were used, these were counted separately. Likewise, studies comparing multiple doses were analysed separately in terms of dose efficacy.

**Table 2.** Data to be extracted for FST systematic review.

| Data to be extracted: |  |
| --- | --- |
| Antidepressant |  |
| Dose | • Dose range (in mg/kg)<br>• Justification of dose (previous publication, pilot study, plasma concentration)<br>• Efficacy of dose (yes, no) |
| Route of administration |  |
| Rodent | • Sex<br>• Strain |
| Treatment | • Duration<br>• Frequency |
| Plasma concentration stated (yes/no) |  |

### Systematic review 2 – A comparison of dose ranges used of fluoxetine and ketamine

#### Search strategy, selection process, inclusion criteria

A systematic review of behavioural animal studies of ketamine and fluoxetine published before 14^th^ February 2020 was conducted via a PubMed search. The following search criteria were used:

*((fluoxetine) OR (ketamine)) AND ((mice) OR (rat)) AND ((antidepressant) OR (depression)) AND ((behavior) or (behaviour)).*

As before, full texts that passed an initial title/abstract screen were examined to identity papers for final inclusion (**Figure 2**), according to the full text inclusion criteria shown in **Table 3**.

**Figure 2.**
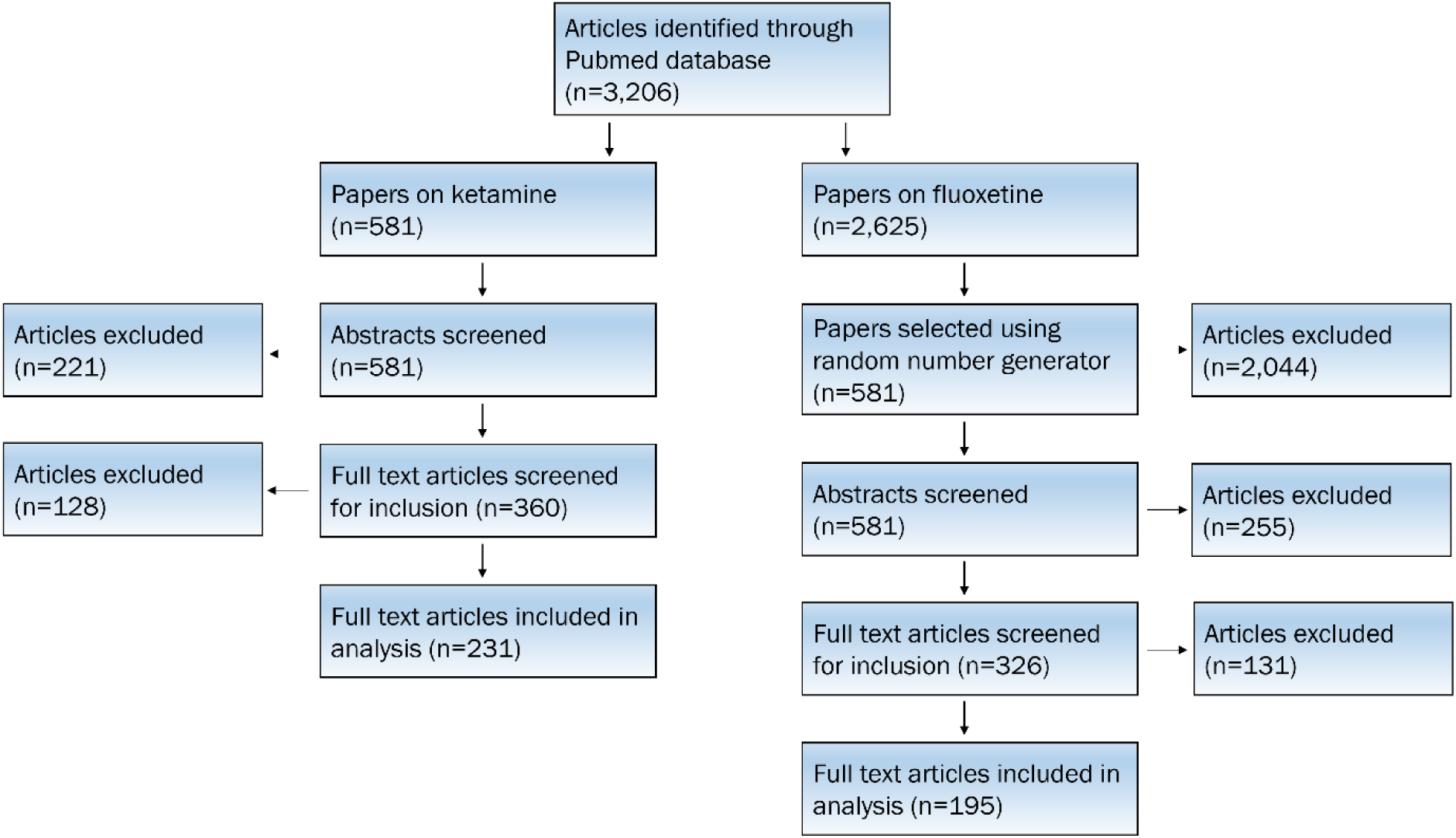
Selection of studies for inclusion in fluoxetine vs ketamine systematic review.

**Table 3.** Full text inclusion criteria for fluoxetine vs ketamine systematic review.

| Inclusion criteria: |
| --- |
| • Drug administered systemically<br>• Reported route of administration, dose, and behavioural readout<br>• Not a review or meta-analysis<br>• English language paper<br>• Paper directly accessible from PubMed<br>• Conducted in rats or mice<br>• Examined (racemic) fluoxetine or ketamine<br>• Ketamine and fluoxetine not co-administered<br>• Examined behavioural measure related to emotion |

#### Data extraction and summary

Similar data of interest were extracted as in the systematic review comparing dose ranges in the FST and ABT (**Table 4**). In this systematic review, dose efficacy was defined as a significant improvement in depression-related behaviours in a given behavioural assay. Again, where multiple doses, species, administration methods or dosing regimens were used, these were analysed separately. For both systematic reviews, data were analysed and figures created using GraphPad Prism 9.

**Table 4.**
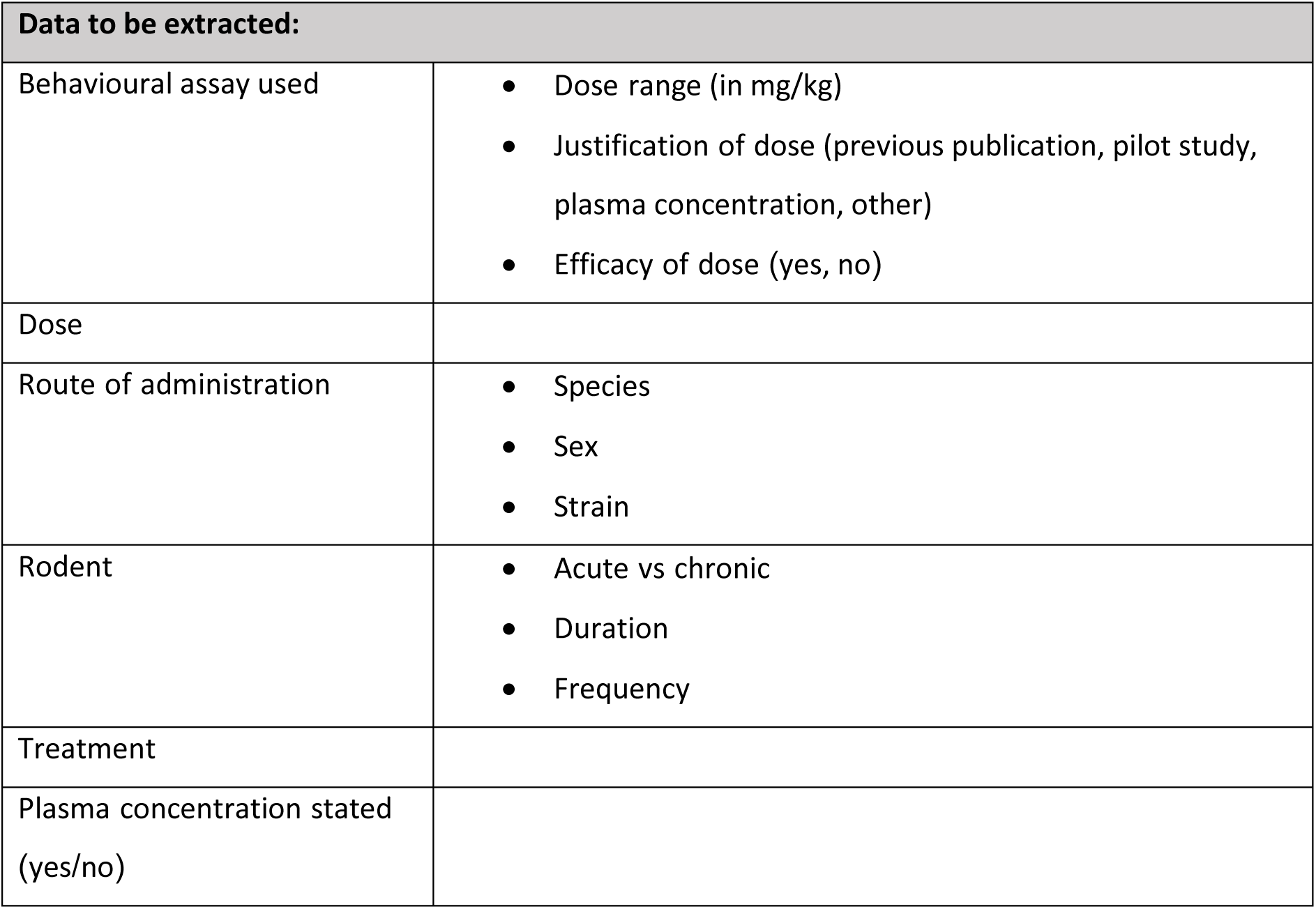
Data to be extracted for fluoxetine vs ketamine systematic review.

## Results

### Systematic review 1 – A comparison of dose ranges used in the Affective Bias Test and Forced Swim Test

The main outcome measure was the range of doses used (**Table 5**) and the rat AED calculated for each drug. Following application of exclusion criteria, 150 papers were included. Raw data can be found in **Data File S1**. 20% of papers studied more than one antidepressant of interest. Most studies in the FST and ABT used male rats and an intraperitoneal route of administration. Full details of the strain and sex of subjects and route of drug administration are detailed in the supplementary materials in **Figure S1**.

**Table 5.** Dose ranges of antidepressants used in ABT and FST studies.

| Drug | N of FST studies | Human clinical dose (mg) | Calculated rat dose (mg/kg) | ABT dose (mg/kg) |  |  | FST dose (mg/kg) |  |  |  |
| --- | --- | --- | --- | --- | --- | --- | --- | --- | --- | --- |
|  |  |  |  | Median | Range | Minimum Effective Dose | Median | Mode | Range | Minimum Effective Dose |
| Agomelatine | 1 | 25 | 1.93 | 0.3 | 0.1–1 | 1 | 10 | NA | 2–50 | 2 |
| Citalopram | 26 | 20 | 1.54 | 0.65 | 0.1–3 | 1 | 10 | 10 | 0.4–60 | 0.4 |
| Clomipramine | 14 | 10 | 0.77 | 1 | 1 | 1 | 20 | 50 | 2.5–60 | 10 |
| Duloxetine | 8 | 60 | 4.63 | 3 | 1–10 | 1 | 15 | 40 | 2.7–80 | 2.7 |
| Fluoxetine | 39 | 20 | 1.54 | 1 | 0.3–3 | 0.3 | 10 | 10 | 1–120 | 3.2 |
| Ketamine* | 58 | 40 | 3.08 | 2 | 1–3 | No effect | 10 | 10 | 0.25–80 | 0.5 |
| Mirtazapine | 5 | 30 | 2.31 | 0.3 | 0.3 | 0.3 | 10 | 5 | 2.5–40 | 2.5 |
| Reboxetine | 16 | 4 | 0.31 | 0.3 | 0.1–1 | 0.1 | 7.8 | 10 | 0.16–128 | 0.16 |
| Sertraline | 9 | 50 | 3.85 | 3 | 0.1–1 | 1 | 10 | 10 | 0.0098–40 | 0.02 |
| Venlafaxine* | 7 | 75 | 5.78 | 3 | 1–10 | 1 | 30 | 10 | 5–80 | 10 |
| Vortioxetine | 3 | 10 | 0.77 | 3 | 1–10 | 3 | 2.5 | 2.5 | 2.5–10 | 2.5 |
| **Total** | 150 |  |  | 1 | 0.1–10 |  | 10 | 10 | 0.0098–128 | 0.02 |

Modal values not presented for ABT new learning studies as each dose per drug only occurred once. Total number of FST studies lower than sum of studies of each antidepressant as some studies examined multiple drugs. Human clinical doses taken from the British National Formulary. \*Ketamine shown to be effective at 1mg/kg when tested in the retrieval model of the ABT but venlafaxine at 3mg/kg had no effect (REF Stuart et al., 2015)

For most drugs, median FST doses were between 1.7-6.5x the AEDs, although this discrepancy was greater for clomipramine and reboxetine whose median doses were 26.0x and 25.2x the corresponding AED, respectively. Overall, 92.4% of FST doses exceeded the relevant calculated AED. Examining each drug separately, in every case over around 80% of doses exceeded the relevant AED, except for sertraline where this was slightly lower (64.3%) (**Table S1**). The median dose used across all FST studies was 10mg/kg and exceeded the rat AED (**Table 5**, **Figure 3A**). Only 37.3% of FST papers tested multiple doses. The use of a large range of doses was also apparent in the FST data. This was greatest for reboxetine which ranged from 0.16–128 mg/kg. In clinical dosing, the range of doses that a clinical can prescribe is quite narrow, in line with the limited therapeutic window of most drugs. In most cases, clinical dose ranges vary by no more than a ½ log unit whereas FST doses for one drug could vary by up to three orders of magnitude.

**Figure 3.**
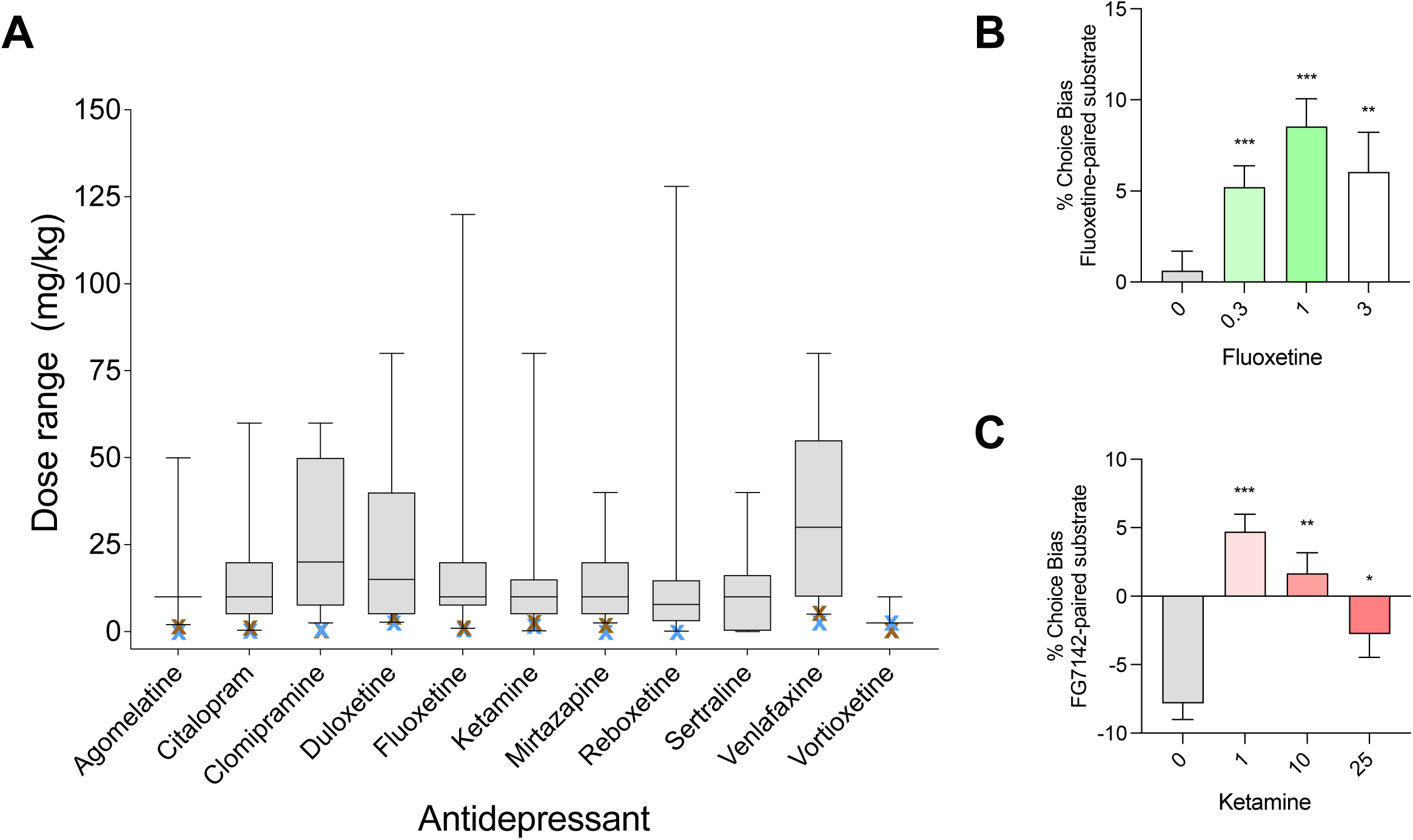
(**A**) Dose ranges used in FST studies. Brown crosses represent rat AEDs for each antidepressant. Blue crosses represent the median dose used in ABT studies of the same drug (**B**) Data from dose-response study of fluoxetine in the ABT (new learning) [25]. (**C**) Data from dose-response study of ketamine in the ABT (24 hour retrieval) [20].

In contrast to the FST, median antidepressant doses for most drugs in the ABT (except duloxetine and vortioxetine) did not exceed the calculated rat AED (**Table 5**, **Figure 3A**). Effects in the ABT were dose-dependent, as shown in **Figure 3B,C**. Above the optimal dose of 1mg/kg, the positive bias toward the fluoxetine-paired substrate begins to diminish (**Figure 3B**). The RAAD ketamine has been shown to modulate affective biases associated with past experiences with an optimal dose 10-fold lower (1 mg/kg) than the average FST dose (and not exceeding the calculated AED for ketamine) (**Figure 4C**). The efficacy of ketamine reduced at higher doses due to generalised amnesic effects [20]. The efficacy of lower doses in the ABT suggests this assay may involve clinically relevant behavioural modulation via underlying mechanisms sensitive to antidepressant drugs at equivalent clinical doses. In line with this, drugs that have given false positives in the FST such as aprepritant and rimonabant do not show efficacy in the ABT [25], with rimonabant inducing a negative bias consistent with its associated depression risk in humans [26].

**Figure 4.**
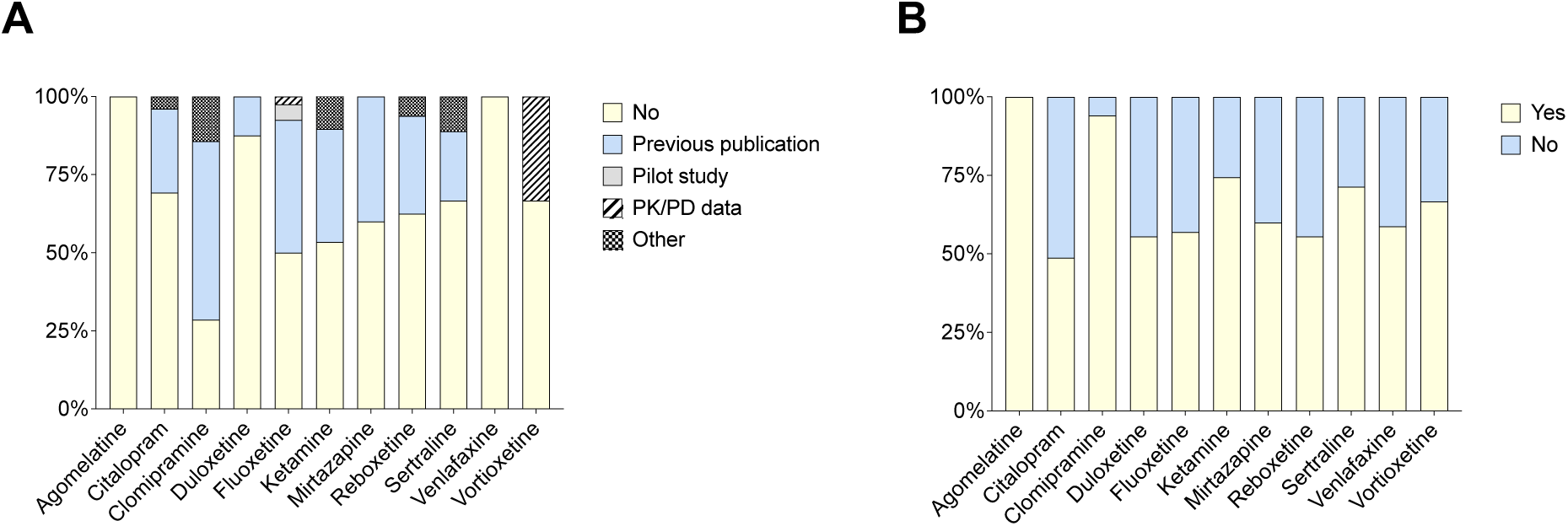
(**A**) Justifications of doses and (**B**) efficacy of doses used in FST papers.

A rationale for the dose chosen in a preclinical study was rarely included or included only in reference to a previous paper where a significant effect was reported. Two of the ABT papers explicitly justified their doses on the basis of pharmacokinetic or pharmacodynamic data (e.g. ED_50_ data from displacement of [^3^H]-citalopram, RO, inhibition constants, brain concentrations). Only 55.1% of FST studies justified their dose choices (**Figure 4A**) and the majority (35.1%) justified their dose based on doses used in previous papers. 4.5% of FST papers gave justifications based on pilot studies and another 4.5% based on other factors. Only one paper referred to pharmacokinetic or pharmacodynamic data (brain SERT occupancy, plasma levels) when justifying their dose choice. All antidepressants showed efficacy in the ABT at one or more dose (**Figure 3C**). 61.2% of doses administered across FST studies exhibited efficacy (**Figure 4B**). The minimum effective doses of most antidepressants in the FST largely matched the relevant AED, indicating that dose efficacy could be seen at more clinically relevant doses.

### Systematic review 2: A comparison of dose ranges used in studies of fluoxetine and ketamine

A total of 581 ketamine and 2,625 fluoxetine papers were identified using our search strategy. Of these, 349 screened ketamine papers were excluded when screened against the inclusion criteria. One more ketamine paper was excluded for using extremely high ketamine doses into the anesthetic range (up to 320 mg/kg), leaving 231 papers in the final analysis. After randomly selecting an equivalent number of fluoxetine papers, 386 papers were excluded and 195 were included in the final analysis. Raw data can be found in **Data File S2**. Full details of subject species and sex are outlined in the supplementary (**Figure S2A,B**). There was a relatively equal split of papers in mice and rats, however most studies used male animals.

The median dose used was 10mg/kg across studies of both drugs exceeding the calculated AEDs for fluoxetine and ketamine by 3.2-6.5x and 1.6-3.2x, respectively (**Table 6**, **Figure 5**). 92.9% of fluoxetine and 65.5% of ketamine doses exceeded the mouse AED while 98.0% of fluoxetine and 76.4% of ketamine doses exceeded the rat AED. The range of doses and maximum dose were considerably larger for ketamine than fluoxetine. Only 18.8% of fluoxetine and 31.0% of ketamine studies tested multiple doses. The frequent use of high doses in these studies likely resulted in plasma concentrations far in excess of what would occur in human patients. **Tables 7** and **8** summarise findings from pharmacokinetic studies in humans and rodents of fluoxetine and ketamine, respectively. These data were found in a literature search and includes plasma concentrations, as well as half-life, clearance and volume of distribution where available. As shown in **Table 7**, an acute oral 20mg dose of fluoxetine in humans elicits lower peak plasma concentrations (11.754-19.1 ng/ml) than an acute 10mg/kg oral dose in rats (61.87 ng/ml). In chronic dosing studies, a highly common 10mg/kg/day i.p. dose in mice resulted in plasma concentrations over an order of magnitude higher (1835.5 ng/ml) than a 20mg/day oral dose in humans (88.8-112.6 ng/ml). Similarly, peak plasma concentrations elicited by an acute clinical 0.5mg/kg intravenous dose of ketamine (133-204 ng/ml) are much lower than are elicited by 10mg/kg and 15mg/kg acute intraperitoneal doses in mice (561.89 ng/ml) (**Table 8**).

**Table 6.** Dose ranges used in ketamine and fluoxetine papers.

| Drug | N of studies | Human clinical dose (mg) | Calculated mouse dose (mg/kg) | Calculated rat dose (mg/kg) | Dose (mg/kg) |  |  |  |
| --- | --- | --- | --- | --- | --- | --- | --- | --- |
|  |  |  |  |  | Median | Mode | Range | Minimum Effective Dose |
| Fluoxetine | 195 | 40 | 3.08 | 1.54 | 10 | 10 | 0.3–80 | 1.25 |
| Ketamine | 232 | 20 | 6.17 | 3.08 | 10 | 10 | 0.1–160 | 0.1 |
| **Total** | 427 |  |  |  | 10 | 10 | 0.1–320 | 0.1 |

**Figure 5.**
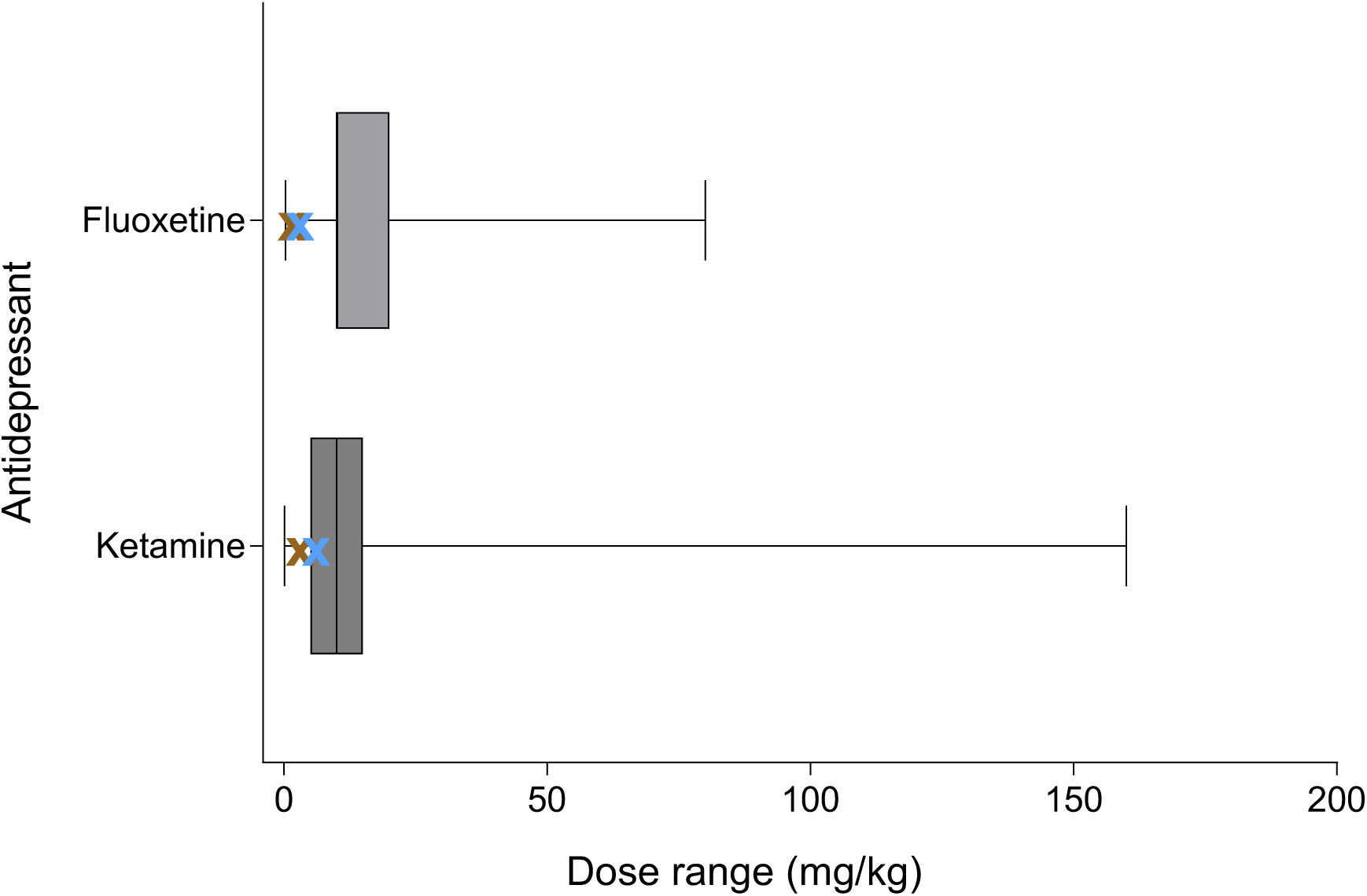
Dose ranges used in ketamine and fluoxetine papers. Blue crosses represent calculated mouse AEDs. Brown crosses represent calculated rat AEDs.

**Table 7.**
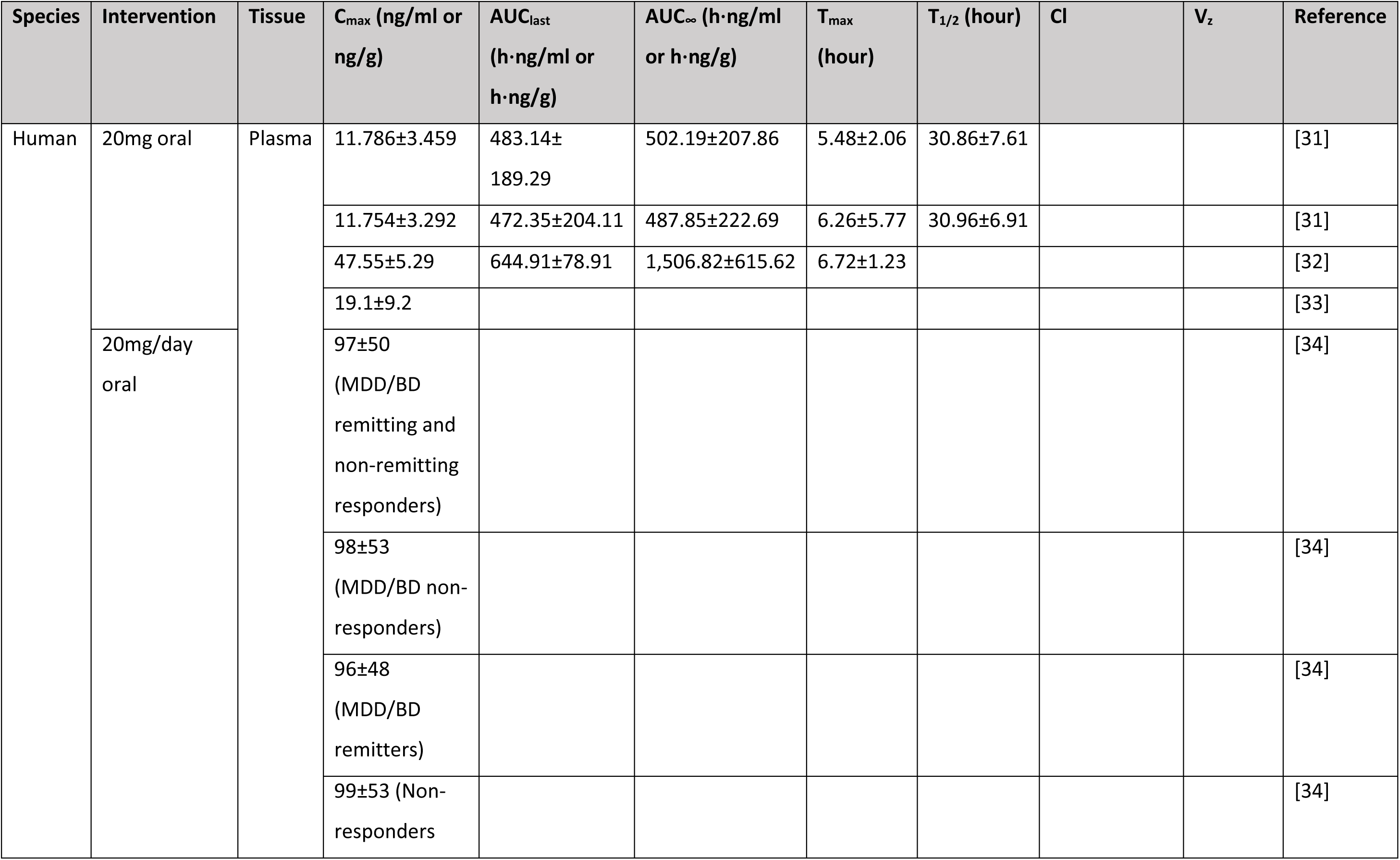
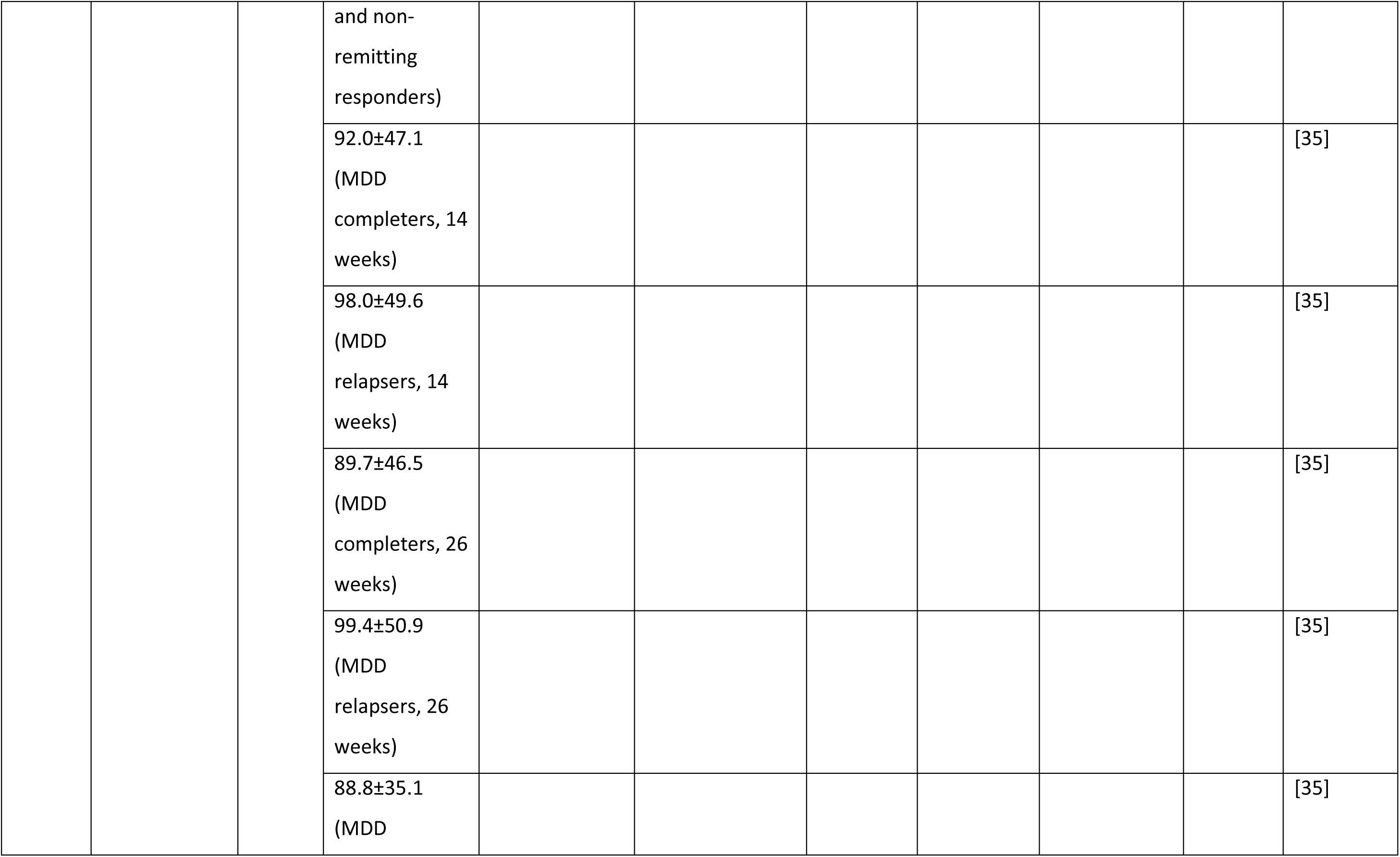
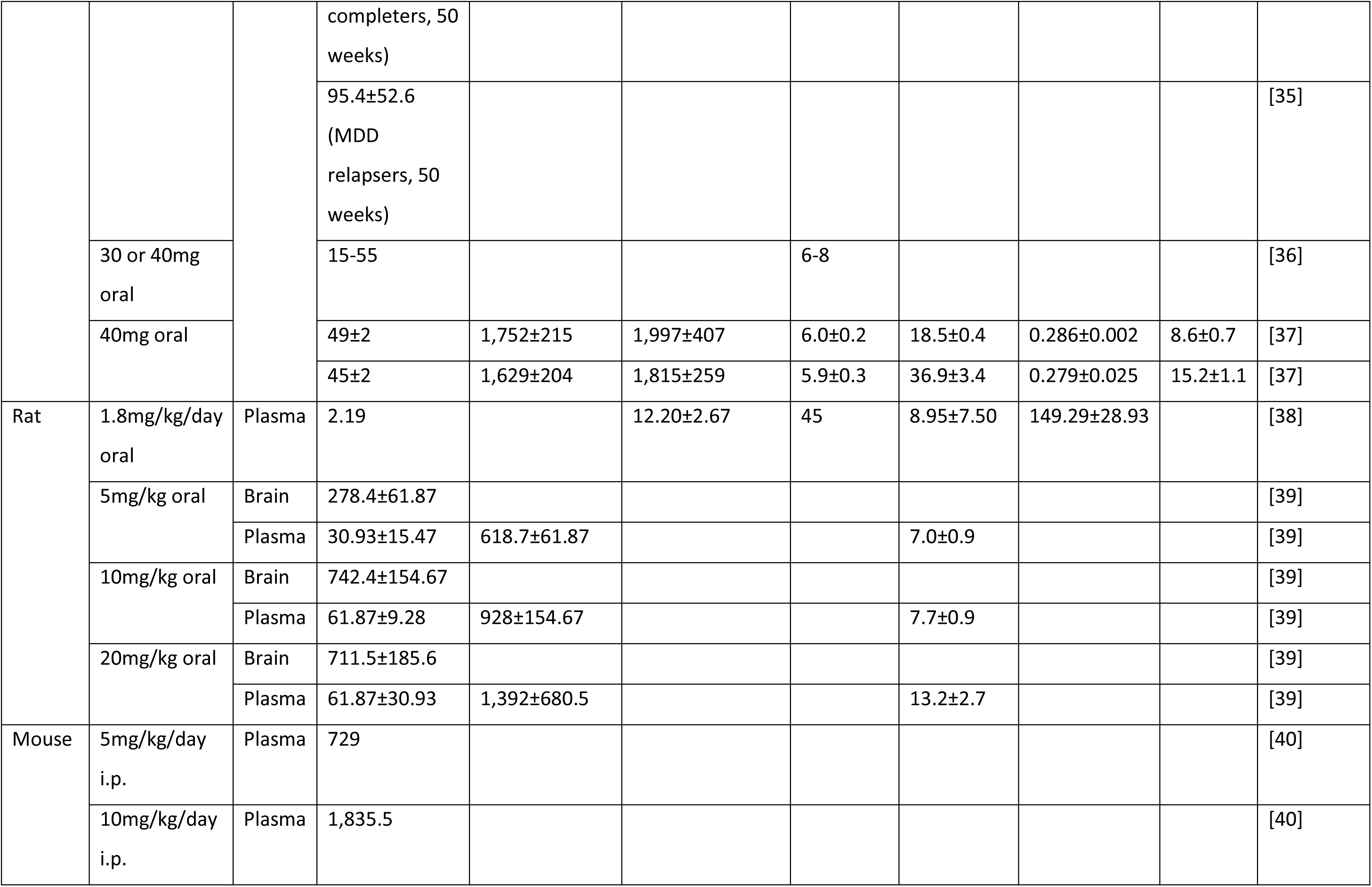
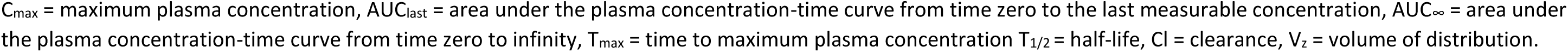
Pharmacokinetics of fluoxetine in humans, rats, and mice.

**Table 8.** Pharmacokinetics of ketamine in humans, rats, and mice.

| Species | Intervention | Tissue | C<sub>max</sub> (ng/ml or ng/g) | AUC<sub>last</sub> (h·ng/ml or h·ng/g) | AUC<sub>∞</sub> (h·ng/ml or h·ng/g) | T<sub>max</sub> (min) | T<sub>1/2</sub> (min) | Cl | V<sub>z</sub> | Reference |
| --- | --- | --- | --- | --- | --- | --- | --- | --- | --- | --- |
| Human | 0.5mg/kg i.v. infusion over 40 mins | Plasma | 177.23±53.8 (BD) | 975.4 (BD) |  |  |  |  |  | [41] |
|  |  |  | 204.13±101.46 (MDD) | 873.5 (MDD) |  |  |  |  |  | [41] |
|  |  |  | 146±16.2 (males) |  | 27,759±2,763 (males) |  | 318±7 (males) | 1,727±132 (males) | 800,955±72,113 (males) | [42] |
|  |  |  | 133±7.2 (females) |  | 23,990±901 (females) |  | 348±17 (females) | 1,475±66 (females) | 732,678±50,344 (females) | [42] |
|  | 10mg i.v. infusion over 30 mins | Plasma | 128.1 |  | 255 |  | 270 | 327 | 39.2 | [43] |
|  | 25mg sublingual tablet |  | 71.1 |  | 184.6 | 45 | 204 | 327 | 39.2 | [43] |
| Rat | 1mg/kg s.c. | Plasma | 76 |  |  |  |  |  |  | [44] |
|  | 3mg/kg s.c. | Plasma | 235 |  |  |  |  |  |  | [44] |
|  | 5mg/kg s.c. | Plasma | 396 |  |  |  |  |  |  | [44] |
|  | 15mg/kg oral | Plasma | 109.11 |  | 90.610 | 5 | 2.5 |  |  | [45] |
|  |  | Brain | 61.77 |  | 67.678 | 5 | 2.35 |  |  | [45] |
|  | 15mg/kg i.p. | Plasma | 524.58 |  | 217.661 | 5 | 15 |  |  | [45] |
|  |  | Brain | 354.06 |  | 179.860 | 5 | 2.55 |  |  | [45] |
|  | 15mg/kg i.n. | Plasma | 149.29 |  | 113.089 | 15 | 2.85 |  |  | [45] |
|  |  | Brain | 77.81 |  | 48.213 | 15 | 2.6 |  |  | [45] |
|  | 40mg/kg i.v. | Plasma | 8,206±697 | 11,410±5,064 |  |  |  |  |  | [41] |
|  | 40mg/kg i.p. | Brain | 32,600±1,400 |  |  |  |  |  |  | [46] |
|  |  | Plasma | 4,738±765 |  | 5,382±2,257 | 6.6±3 | 612±312 |  |  | [47] |
|  | 40mg/kg/day i.p. | Plasma | 6,365±820 |  | 5,749±513 | 10.2±0 | 168±78 |  |  | [47] |
| Mouse | 10mg/kg i.p. | Plasma | 561.89±86.09 | 177.9 |  |  |  |  |  | [41] |
|  |  | Brain | 1,162.34±202.05 | 492.2 |  |  |  |  |  | [41] |

C<sub>max</sub> = maximum plasma concentration, AUC<sub>last</sub> = area under the plasma concentration-time curve from time zero to the last measurable concentration, AUC<sub>∞</sub> = area under the plasma concentration-time curve from time zero to infinity, T<sub>max</sub> = time to maximum plasma concentration T<sub>1/2</sub> = half-life, Cl = clearance, V<sub>z</sub> = volume of distribution.

As plasma concentrations are directly related to receptor occupancy, this suggests that typical doses used in rodent preclinical studies may elicit higher receptor occupancies than are achieved with clinical doses. A 20mg dose of fluoxetine in humans results in serotonin transporter (SERT) receptor occupancy of around 75% [48] whereas a typical 10mg/kg oral dose in rats results in over 95% SERT receptor occupancy [49]. Data regarding N-methyl-D-aspartate (NMDA) receptor occupancy in humans is lacking, however Shaffer et al calculated predicted NMDA receptor occupancy across several species based on unbound ketamine plasma concentrations [50]. Predicted NMDA receptor occupancy in humans following a 0.5mg/kg intravenous infusion elicited similar maximal mean receptor occupancies (∼30%) as a 10mg/kg intraperitoneal dose in rats, but RO levels differed over time between species. This indicates that standard rodent doses produce different absolute receptor occupancy and/or receptor occupancy-time relationships than clinical doses of fluoxetine and ketamine. At high doses, it is also difficult to disentangle target and off-target receptor effects in a behavioural assay, as shown in **Tables S2** and **S3**, which present the binding affinities of fluoxetine and ketamine to various receptors in rodents and humans, demonstrating how they may engage off-target receptors at higher concentrations.

The greatest proportion of papers administered drugs intraperitoneally (fluoxetine: 58.3%, ketamine: 95.2%), however a significant proportion of fluoxetine studies (37.1%) utilised oral dosing methods (**Figure S2C**), consistent with its oral administration in humans. Ketamine’s antidepressant utility has mainly been examined in humans through intravenous intervention [51], however the vast majority of ketamine studies (95.3%) in this review utilised intraperitoneal dosing methods. Most (62.1%) fluoxetine papers utilised a chronic dosing scheme, consistent with its delayed onset and need for daily administration in humans [15] (**Figure S2D**). In contrast, the antidepressant effects of ketamine emerge almost immediately after a single dose [52], thus justifying the use of acute dosing regimens in ketamine studies in this review (88.8%). Few papers combined both (fluoxetine: 6.2%, ketamine: 2.7%).

Most studies reviewed here did not justify their dose selection (fluoxetine: 64.1%, ketamine: 52.0%) (**Figure 6A**). Where a justification was given, again the most common justification given was that dose(s) had been used in previous studies (fluoxetine: 29.2%, ketamine: 41.5%), with far fewer papers justifying their dose selection on pilot studies (fluoxetine: 2.6%, ketamine: 1.3%) or other factors (fluoxetine: 1.5%, ketamine: 4.8%). Four fluoxetine papers justified their dose choices based on typical brain or plasma concentrations elicited therapeutically but only one ketamine paper justified dose choice based on pharmacokinetic or pharmacodynamic data, doing so based on NMDA receptor occupancy. As shown in **Figure 6B**, most doses administered (fluoxetine: 86.6%, ketamine: 78.8%) evoked a significant effect in the given behavioural assay and, again the minimum effective dose for fluoxetine matched the AED and was even lower than the AED for ketamine.

**Figure 6.**
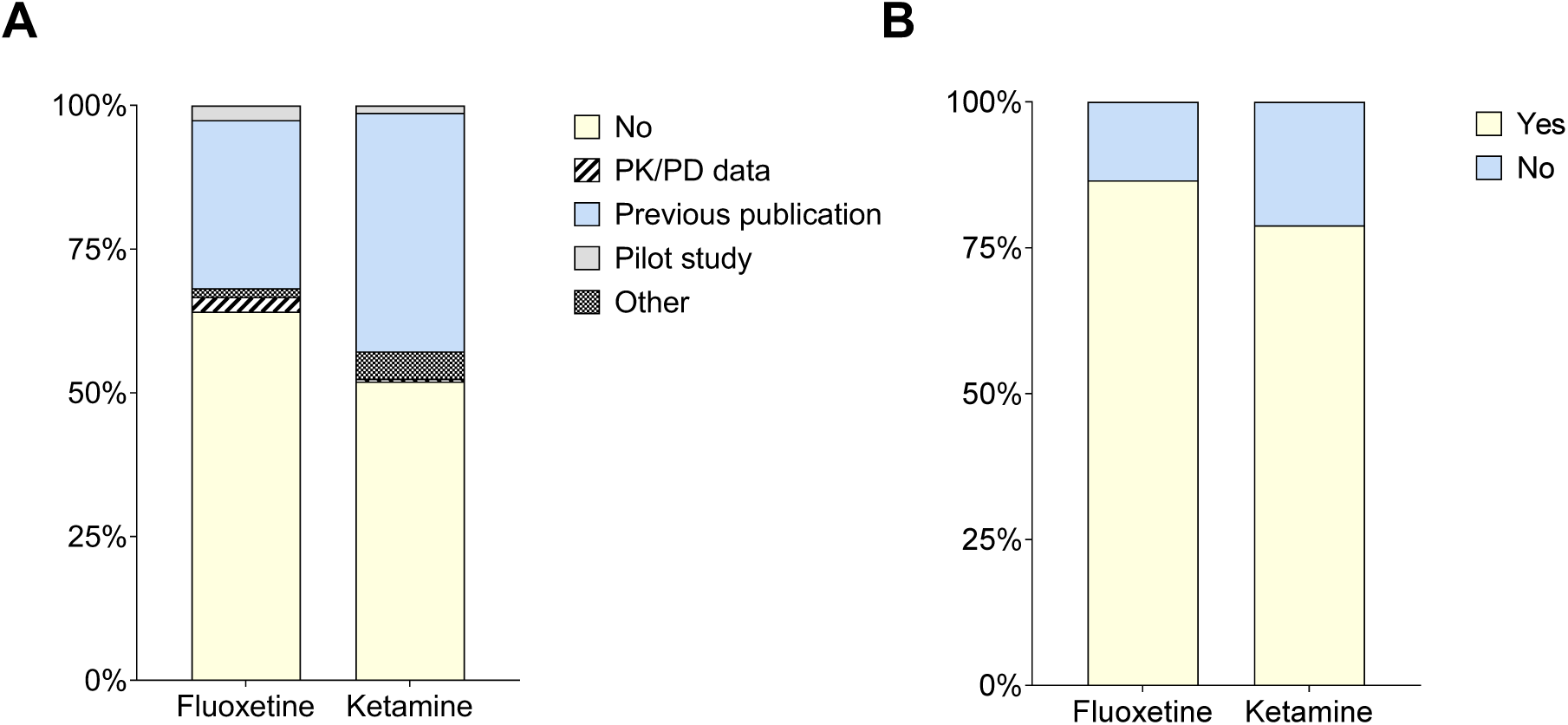
(**A**) Justifications of doses and (**B**) efficacy of doses used in ketamine and fluoxetine papers.

## Discussion

Both systematic reviews found evidence that doses of antidepressant drugs used in pre-clinical research involving traditional models of depression exceed the AED by several orders of magnitude, potentially limiting the translational relevance of the arising data. In contrast to studies in the FST, doses which were most effective in the ABT aligned more closely with clinical doses and associated receptor occupancy. Where studies used doses which led to plasma levels exceeding those found clinically, interpretation of the arising behavioural effects may be confounded by effects on off-target receptors or exceeding clinically relevant levels of receptor occupancy. Achieving clinically relevant levels of receptor occupancy using appropriate doses in preclinical models is critical for translational validity and to increase the predictive value of the preclinical data. Even the more selective second generation antidepressants will produce very different pharmacodynamic profiles when administered at higher doses [53]. This is illustrated in **Figure 7A** for fluoxetine, where increasing doses lead to higher occupancy not only at SERT but also at off-target receptors such as the 5-HT2C and 5-HT2A receptors. A clear clinical example highlighting the importance of dose on the behavioural effects of a drug is seen with ketamine. At low doses (0.5 mg/kg per 40 min i.v. or peak plasma concentrations of 70-200 ng/ml [54]) ketamine demonstrates rapid antidepressant efficacy [24], moderate doses (0.1-0.3 mg/kg [55]) are analgesic, and high doses (1-4.5 mg/kg over 60 seconds or peak plasma concentrations of 2000-3000 ng/ml [54]) are anaesthetic. As shown in **Figure 7B**, as the dose of ketamine increases, so does its occupancy of not only the NMDA receptor but also off-target receptors like Mu2 opioid and delta opioid receptors. Indeed, there is evidence that opioid receptors are involved in the analgesic effects which are reversed by the opioid receptor antagonist naloxone [56, 57]. This dose-dependent receptor engagement explains the diverse clinical effects of ketamine and highlights the importance of using the appropriate dose in animal models.

**Figure 7.**
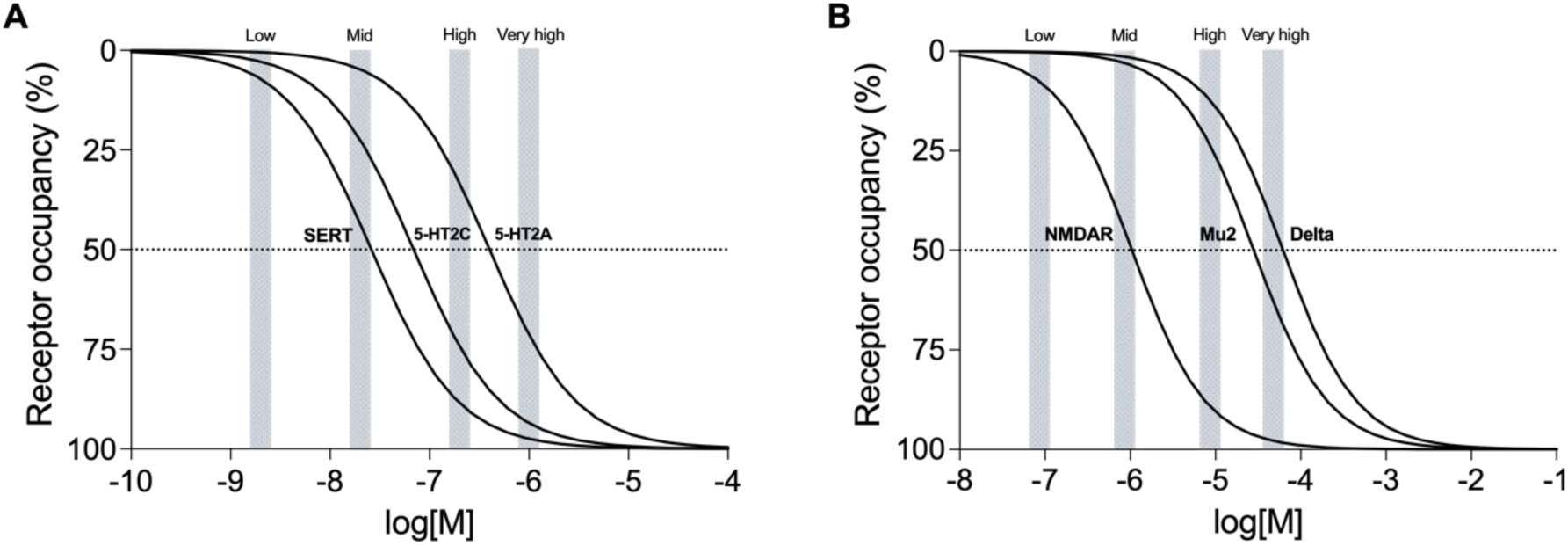
Schematic representation of fluoxetine and ketamine receptor occupancy at three of their molecular targets, with inhibition curves showing rat IC50s for the SERT, 5-HT2C, and 5-HT2A receptors and NMDA, Mu2 opioid and delta opioid receptors for fluoxetine and ketamine, respectively. (**A**) Kis (µM): SERT = 0.026, 5-HT2C = 0.069, 5-HT2A = 0.401. Grey bars represent maximum plasma concentrations elicited at low (0.1 mg/kg), medium (1 mg/kg), high (10 mg/kg), and very high (50 mg/kg) fluoxetine doses. (**B**) Kis (µM): NMDA = 1.048, Mu2 opioid = 28.1, delta opioid = 62. Grey bars represent maximum plasma concentrations elicited at low (0.1 mg/kg), medium (1 mg/kg), high (10 mg/kg), and very high (50 mg/kg) ketamine doses. Plasma concentrations are estimated based on a 40 mg/kg i.v. dose of ketamine eliciting plasma concentrations of ∼8,206 ng/ml in rats 65). Plasma concentrations are estimated based on a 10 mg/kg oral dose of fluoxetine eliciting plasma concentrations of ∼61.87 ng/ml in rats [39]. For both drugs, plasma concentrations for a 10-fold higher dose are assumed to elicit a 10-fold higher plasma concentration.

When drugs are administered to an animal at doses which lead to plasma levels that exceed those achieved clinically, this complicates the conclusions that can be drawn from preclinical studies. Physiological or behavioural effects elicited at high doses may not engage clinically relevant mechanisms and be misinterpreted as predicting clinical efficacy. This risk is increased when high doses are used in combination with animal behavioural assays with weak translational validity, such as assays susceptible to behavioural confounds (e.g. locomotor activity). Considering that most animal models for depression were developed based on their ability to predict a pharmacological effect, one thus cannot be confident that the effects seen in assays such as the FST are generated by clinically relevant occupancy of the target receptor and are therefore translatable to patients. This may lead to inappropriate conclusions, for instance regarding the mechanisms of a known antidepressant and identification of novel drug targets or regarding the potential clinical utility of a novel compound.

An important finding from this review was the consistent lack of pharmacological rationale for dose selection in preclinical studies of antidepressants, with studies most often relying on doses used in previous papers and where significant effects were reported. This has significant implications for study quality and the translational validity of study findings. Without robust justification based on pharmacokinetic or pharmacodynamic data, the chosen doses may not accurately reflect clinically relevant concentrations. Additionally, the assumption that previous publications observing significant effects is a justification for dose selection perpetuates the problem and can undermine the rigor and credibility of the studies. This habitual reliance of preclinical depression studies on convention rather than empirical data contributes to a cycle of poorly justified dosing, which can mislead future research efforts and delay the development of effective antidepressants. The findings from these systematic reviews suggest improving the rigor of dose selection and reporting in preclinical studies could enhance the clinical relevance of research in animal models particular for studies aimed at identifying underlying mechanisms.

One possible factor contributing to the widespread use of high doses may stem from animal studies frequently being underpowered [10]. Low statistical power in studies increases the chance of false negatives and false positives and can result in the overestimation of the true effect size as only those studies that by random chance show large effects will achieve significance [10]. It is widely appreciated that researchers are under tremendous pressure to publish and research that results in a statistically significant finding is more likely to be published than negative data [58]. For this reason, researchers are incentivised to engage in research practices that expedite the publication of their work and may contribute to poor reproducibility and translation [12]. Researchers may administer higher drug doses in underpowered behavioural studies to reduce variation and increase their likelihood of detecting behavioural effects and achieving significant results. This practice may be used to compensate for the difficulty of detecting true effects at lower, clinically relevant doses. To address these issues, studies should prioritise rigorous power calculations based on estimated effect sizes. This approach ensures adequate sample sizes to detect behaviourally relevant and clinically meaningful changes induced by antidepressant treatments, thereby enhancing the reliability and translational potential of preclinical research in psychopharmacology.

## Conclusions and recommendations

These systematic reviews revealed the pervasive use of potentially inappropriate dose ranges in animal behavioural studies of MDD. This widespread use of high doses was evident both across multiple drugs and across multiple behavioural assays. This phenomenon may be contributing to failures in the translation of preclinical findings, with far reaching implications. This study only investigated two antidepressants in MDD research and future research examining dose ranges across different psychiatric drug classes is required to assess whether this phenomenon occurs more generally across the field.

Unfortunately, the need for appropriate dose selection has been known for some time but these reviews suggest the issues remain a common problem in antidepressant research. As found in these studies, the rationale for the choice of dose(s) rarely used pharmacokinetic and pharmacodynamic data. As early as 2000, in a sample of over 4000 animal studies of antipsychotics, found that most papers did not discuss their method of dose selection. More commonly, researchers used the same dose as a previous study and very rarely did researchers justify their dose choice based on clinical relevance [59]. However, as shown in our search of rodent and human receptor occupancy data for 59 psychiatric drugs (**Figure S1**), these data are often not easily available. This highlights the need for improved public access to such data to facilitate dose selection and we suggest the development of an open-access database containing all available industry and academic pharmacodynamic and pharmacokinetic data of psychiatric drugs. Until such a database exists, estimating the dose of a drug to use in animals will remain a challenge for academic researchers. However, these systematic reviews suggest greater consideration of the minimal effective dose, predicted receptor occupancy, effect size and appropriate estimations of sample size could improve the quality of studies in pre-clinical MDD research. It is also interesting to observe that the effective dose in the translational ABT test align much more closely with the AED and clinical doses suggesting improving the clinical relevance of the behavioural readouts used in animal studies can also help address some of these issues.

## Acknowledgements

Sophie Kostine^1^, E Wright^1^, S Harris^1^,

## Supplementary materials and methods

### Additional info for methods and materials

#### Rationale for choice of behavioural tests and drugs included in these systematic reviews

The FST was once widely considered the ‘gold-standard’ test for studying antidepressant efficacy in rodents [1]. In this task, the animal is placed in a container of water from which it cannot escape and the time taken for the animal to switch from an active coping strategy (i.e. swimming, climbing) to a passive coping strategy (i.e. immobility) is measured [2]. In pharmacological studies the FST has shown sensitivity to antidepressant treatment, including TCAs and monoamine oxidase inhibitors (MAOIs) [3]. Later, a modified version of the FST was introduced that provided greater reliability when assessing SSRIs [4]. The validity of the FST as a behavioural assay for depression has been widely criticised [2, 5–7]. The FST has yielded a number of false positives and false negatives [8, 9] and has not produced any novel antidepressants [6]. In a recent retrospective review of 109 compounds tested in the FST, only 28% were investigated in humans and, of those, only seven successfully predicted antidepressant effects in humans [9]. In a recent systematic review of studies that assessed N-methyl-D-aspartate receptor (NMDAR) antagonists in either the FST or the related Tail Suspension Test (TST), immobility time in these tasks was not found to be a good predictor of clinical efficacy in clinical trials [10]. Another criticism is that the FST is often interpreted as modelling ‘depression-like’ behaviour or ‘behavioural despair’ in rodents, especially in mechanistic studies. Such conclusions are a potentially dangerous leap of logic with no empirical basis in the pathophysiology of depression [11, 12]. Furthermore, although the FST is sensitive to stress-related interventions [13], it is not so sensitive to other depression-related risk factors in humans, such as early life adversity [14], thus raising further uncertainty over its translational utility.

An alternative approach is to design tasks based on neuropsychological theories of depression. For instance, modern theories highlight the importance of altered affective processing in the pathogenesis of depression [15, 16]. This concept inspired the development of the ABT which examines how manipulations of emotional state (e.g. by antidepressant drugs) affect reward learning and memory in rodents [16]. Rats independently learn to associate two digging mediums with an equally valued food reward, one of which is paired with an emotional manipulation. A pharmacologically induced affective bias can be identified during a preference test in which rats are presented with both substrates. A bias toward digging in the drug-paired bowl indicates a positive bias whereas a bias away from it indicates a negative bias [17]. Acute treatment with several antidepressants, including SSRIs, noradrenaline reuptake inhibitors (NRIs), serotonin and noradrenaline reuptake inhibitors (SSNRIs), TCAs, and atypical antidepressants, as well as putative non-monoaminergic antidepressants, elicit positive biases in the ABT [17–21]. The ABT also differentiates between delayed and rapid acting antidepressants providing evidence that distinct neuropsychological mechanisms underlie the temporal differences in their clinical effects [21]. Given the differences in terms of translational validity, the FST and ABT were chosen as exemplar behavioural assays to compare in these reviews.

Most currently available antidepressant medications are modelled on early drugs (TCAs) whose antidepressant effects were discovered serendipitously in the 1950s [22]. Relying on the same mechanism of action of enhancing serotonergic function in the brain by increasing synaptic monoamine levels of serotonin [23], modern antidepressants, such as the SSRI fluoxetine which was introduced to clinical practice as a treatment for depression in 1988 [24], exhibit no greater efficacy than early antidepressants. Approximately 30% of patients do not respond to SSRIs [25] and, even id they do, antidepressant effects can take several weeks to develop [15].

This has led some to propose other neurobiological mechanisms underlying depression and shifted focus to novel antidepressant candidates [24]. Interest in a new class of rapid-acting antidepressants was prompted by evidence that a single sub-anaesthetic dose of ketamine, an NMDAR antagonist, results in antidepressant effects in humans within 72 hours [26]. It was subsequently shown that ketamine demonstrates clinical efficacy within hours, even among treatment-resistant populations [27]. Furthermore, ketamine’s effects seem to outlast its pharmacokinetics, with many patients demonstrating clinical improvements lasting up 6 weeks [28], despite its relatively short half-life of only 2-4 hours [29]. Given that fluoxetine is the most extensively prescribed antidepressant and that the s-enantiomer of ketamine (esketamine) is the first rapid-acting antidepressant to be approved in the UK (in 2020) and the US (in 2019), fluoxetine and ketamine drugs were chosen as exemplar delayed-onset and RAADs to compare in systematic review 2.

### Additional info for results

Regarding the characteristics of subjects used in each task in systematic review 1, two of the ABT studies used male Lister-hooded rats while one used male Sprague Dawley and another Sprague Dawley rats from both sexes. The majority of FST papers used Wistar (48.7%) or Sprague Dawley (44.2%) rats, with far fewer using albino (1.3%), Flinders Sensitive Line (3.2%), Roman (1.3%), and Wistar Kyoto (1.3%) rats. This is shown separated by drug in **Figure S1A**. Most (89.4%) of FST papers used male rats, with 2.6% using female rats and 4.6% using both sexes, as shown by drug in **Figure S1B**. 3.3% did not report the sex of the animals used.

**Figure S1.**
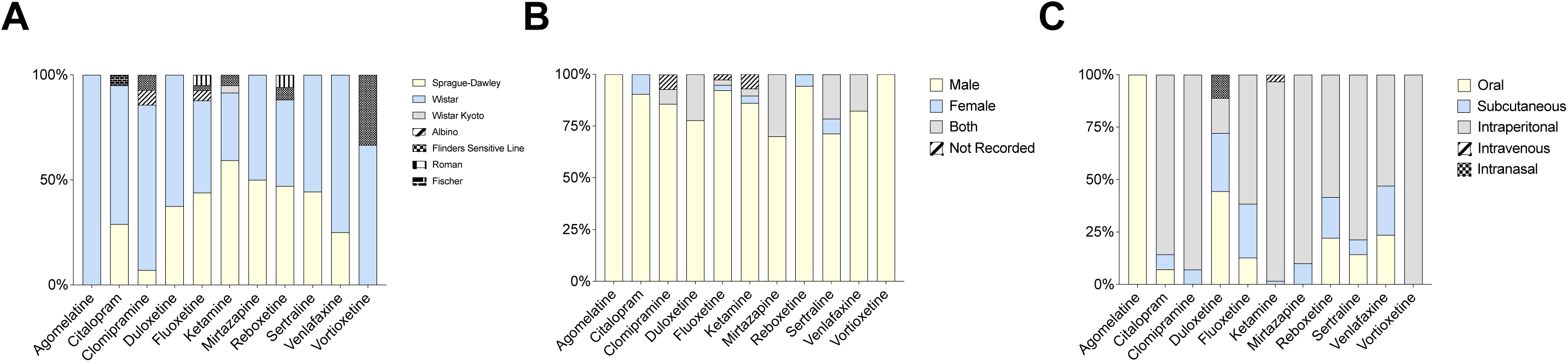
(**A**) Rodent strain, (**B**) rodent sex, and (**C**) routes of administration used in FST papers.

**Table S1.** Proportion of doses exceeding the relevant calculated rat AED, separated by dose.

| Drug | % doses exceeding rat AED |
| --- | --- |
| Agomelatine | 100.0% |
| Citalopram | 88.1% |
| Clomipramine | 100.0% |
| Duloxetine | 88.9% |
| Fluoxetine | 96.9% |
| Ketamine | 83.3% |
| Mirtazapine | 100.0% |
| Reboxetine | 94.4% |
| Sertraline | 64.3% |
| Venlafaxine | 94.1% |
| Vortioxetine | 100.0% |
| **Total** | 89.9% |

In systematic review 2, just over half of fluoxetine (61.0%) and ketamine (ketamine 55.1%) papers used mice, with slightly fewer using rats (fluoxetine: 38.5%, ketamine: 42.7%), and a minority using both (fluoxetine: 0.5%, ketamine: 2.3%) (**Figure S2A**). The majority of fluoxetine (78.0%) and ketamine (71.1%) papers used male animals, with far fewer using female animals (fluoxetine: 10.3%, ketamine: 8.6%), or a combination of both (fluoxetine: 4.1%, ketamine: 3.5%) (**Figure S2B**). 7.7% of fluoxetine papers and 16.8% of ketamine did not record subjects’ sex.

**Figure S2.**
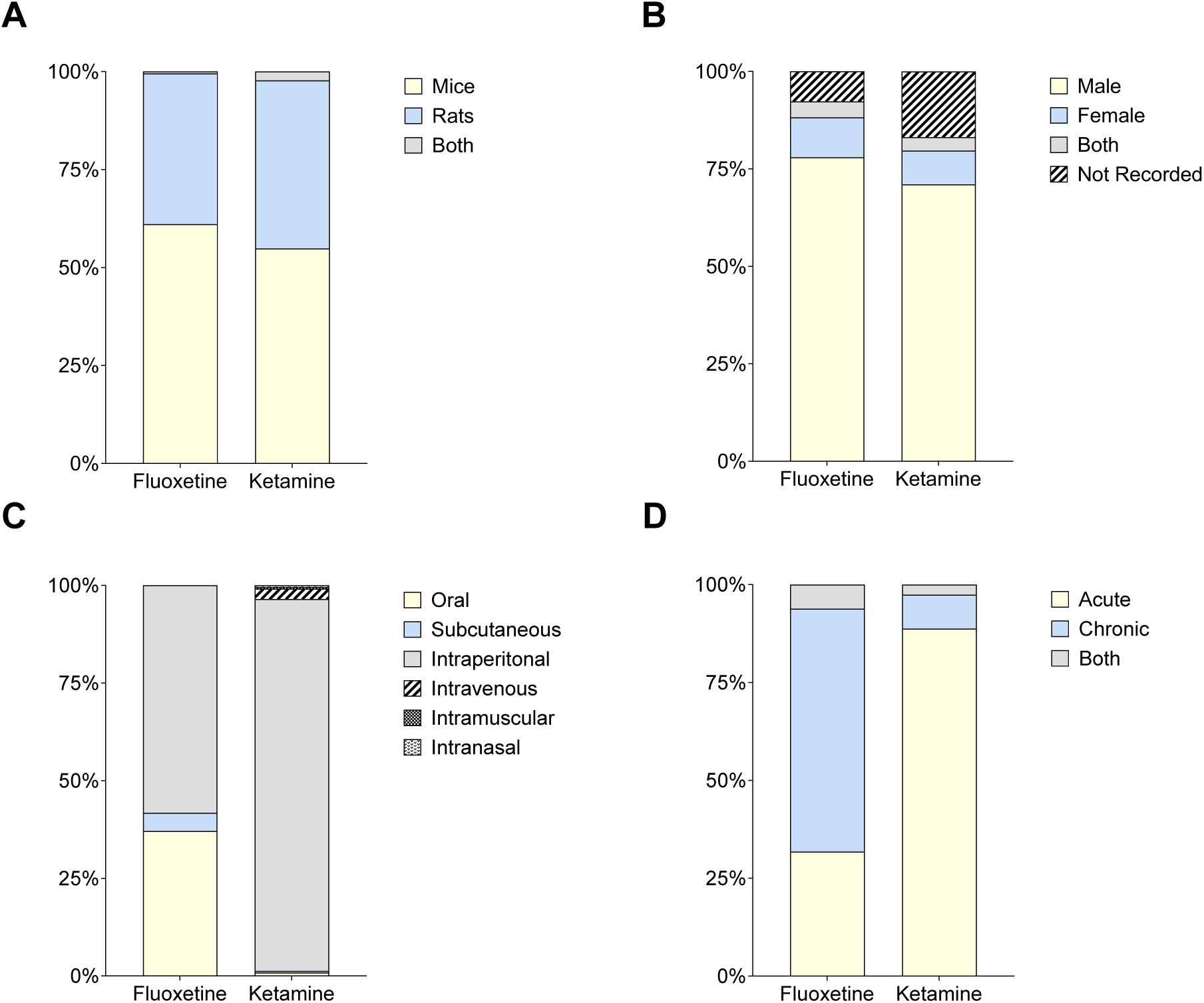
(**A**) Rodent species, (**B**) rodent sex, (**C**) routes of administration, and (**D**) treatment regimes used in ketamine and fluoxetine papers.

Data for fluoxetine binding affinities below were taken from the PDSP database [30], and ketamine affinities were taken from this database as well as a paper by Zanos and colleagues [31]. Besides SERT, fluoxetine demonstrates some affinity at several serotonin receptors, muscarinic receptors, adrenergic receptors, and histamine H1 receptor (**Table S2**). Off-target activation of these receptors by TCAs is known to cause side effects like dizziness, memory problems and tiredness [32]. Likewise, ketamine exhibits some affinity for several receptors other than the NMDA receptor, including the D_2_ and sigma-1/2 receptors (**Table S3**). At high doses, engagement of these receptors is more likely and could bring about unintended behavioural and/or physiological effects that are misconstrued as indicating depression-relevant efficacy.

**Table S2.** Binding affinities of fluoxetine to its receptor targets.

| Receptor target | Species | Ki (μM) | Reference |
| --- | --- | --- | --- |
| **SERT** | Rat | 0.026 | [33-42] |
|  | Human | 0.025 | [37, 42-48] |
| **5-HT1A** | Rat | 8.313 | [37] |
| **5-HT1B** | Rat | 6.166 | [49] |
| **5-HT2** | Rat | 1.410 | [34, 49] |
|  | Human | 0.28 | [50] |
| **5-HT2A** | Rat | 0.401 | [37, 39, 51-53] |
|  | Human | 0.197 | [53, 54] |
| **5-HT2B** | Human | 5.03 | [52] |
| **5-HT2C** | Rat | 0.069 | [39, 51-53] |
|  | Human | 0.315 | [45, 53-55] |
| **5-HT6** | Rat | 1.77 | [56] |
| **5-HT7** | Rat | 1 | [57] |
| **Adrenergic alpha 1** | Rat | 2.491 | [34, 40] |
|  | Human | 3.556 | [37, 43, 50, 58] |
| **Adrenergic alpha 1A** | Human | 2 | [55] |
| **Adrenergic alpha 2** | Rat | 3.906 | [40, 51] |
|  | Human | 2.19 | [37, 58] |
| **Cholinergic muscarinic** | Rat | 1.806 | [37, 51] |
|  | Human | 1.295 | [50, 58] |
| **DAT** | Rat | 3.097 | [34, 35, 59, 60] |
|  | Human | 3.446 | [43, 44, 46, 47] |
| **Histamine H1** | Rat | 3.2 | [51] |
|  | Human | 3.975 | [50, 58, 61] |
| **NET** | Rat | 0.716 | [34-37, 40, 42, 60, 62] |
|  | Human | 0.818 | [37, 43, 44, 46, 47] |
| **Sigma 1** | Rat | 1.911 | [30] |
| **Sigma 2** | Rat | 1.121 | [30] |

Binding affinities for fluoxetine were taken from the PDSP database [30], accessed 24<sup>th</sup> July 2024. Only studies where the tissue source was defined were excluded. Where multiple papers have been published for the same receptor in the same species, average values are stated.

**Table S3.** Binding affinities of ketamine to its receptor targets.

| Receptor target | Species | K<sub>i</sub> (μM) | Reference |
| --- | --- | --- | --- |
| **NMDAR** | Rat | 1.048 | [63-79] |
|  | Mouse | 0.58 | [70] |
|  | Human | 1.101 | [77, 80, 81] |
| **5-HT2** | Rat | 15 | [82] |
| **5-HT3** | Mouse | 96.9 | [82] |
| **Cholinergic muscarinic M1** | Human | 45 | [83] |
| **Cholinergic muscarinic M2** | Human | 294 | [83] |
| **Cholinergic muscarinic M3** | Human | 246 | [83] |
| **D2** | Rat | 1 | [73] |
|  | Human | 0.525 | [73, 74, 76] |
| **DAT** | Rat | 62.9 | [84] |
| **Delta opioid** | Human | 272 | [85] |
| **Kappa opioid** | Human | 26.55 | [85, 86] |
| **Mu opioid** | Human | 42.1 | [85] |
| **Sigma opioid** | Rat | 0.946 | [87] |
| **Sigma 1/2** | Rat | 5.075 | [71, 88] |
| **Sigma 1** | Rat | 139.6 | [89] |
| **Sigma 2** | Rat | 26.3 | [89] |

Binding affinities for ketamine were taken from the PDSP database [30], accessed 24<sup>th</sup> July 2024, as well as a paper by Zanos and colleagues [31]. Only studies of racemic ketamine and where the tissue source was defined were included. Kis over 10,000 are not included.

### Systematic review 3 – Receptor occupancy data for psychiatric drugs

#### Methods

##### Search strategy, selection process, inclusion criteria

A PubMed search was conducted of all rodent and human receptor occupancy studies of psychiatric drugs. The following search criteria were used:

*(DRUG) AND (occupancy) AND ((rat) OR (mouse)) OR (human)*

with a separate search conducted for each of the following 59 psychiatric drugs: alprazolam, amisulpride, aripiprazole, asenapine, benperidol, bromazepam, bromperidol, buspirone, chlordiazepoxide, chlorpothixene, clobazam, clonazepam, clonidine, clorazepate, clozapine, dexamfetamine, divalproex sodium, flupentixol, fluspirilen, gabapentin, guanfacine, haloperidol, hydroxyzine, iloperidone, lamotrigine, levetiracetam, levomepromazine, lisdexamfetamine, lithium salts, lorazepam, lurasidone, melperone, meprobamate, methamphetamine, methylphenidate, modafinil, olanzapine, oxazepam, oxcarbazepine, paliperidone, perazine, perphenazine, pimozide, pregabalin, promethazine, prothipendyl, quetiapine, risperidone, sibutramine, sodium valproate, sulpiride, thioridazine, thiothixene, tofisopam, topiramate, trifluoperazine, ziprasidone, zotepine, zuclopenthixol.

However, it was decided that the search terms used were too restrictive and so a subsequent search with broader search terms was conducted to identify papers published before 10^th^ February 2022:

*(DRUG) (occupancy) (SPECIES)*

with separate searches conduced mice, rats, and humans for the same drugs as listed above (**Figure S3**). Abstracts and full text studies were screened and assessed against the inclusion criteria listed in **Table S4**.

**Figure S3.**
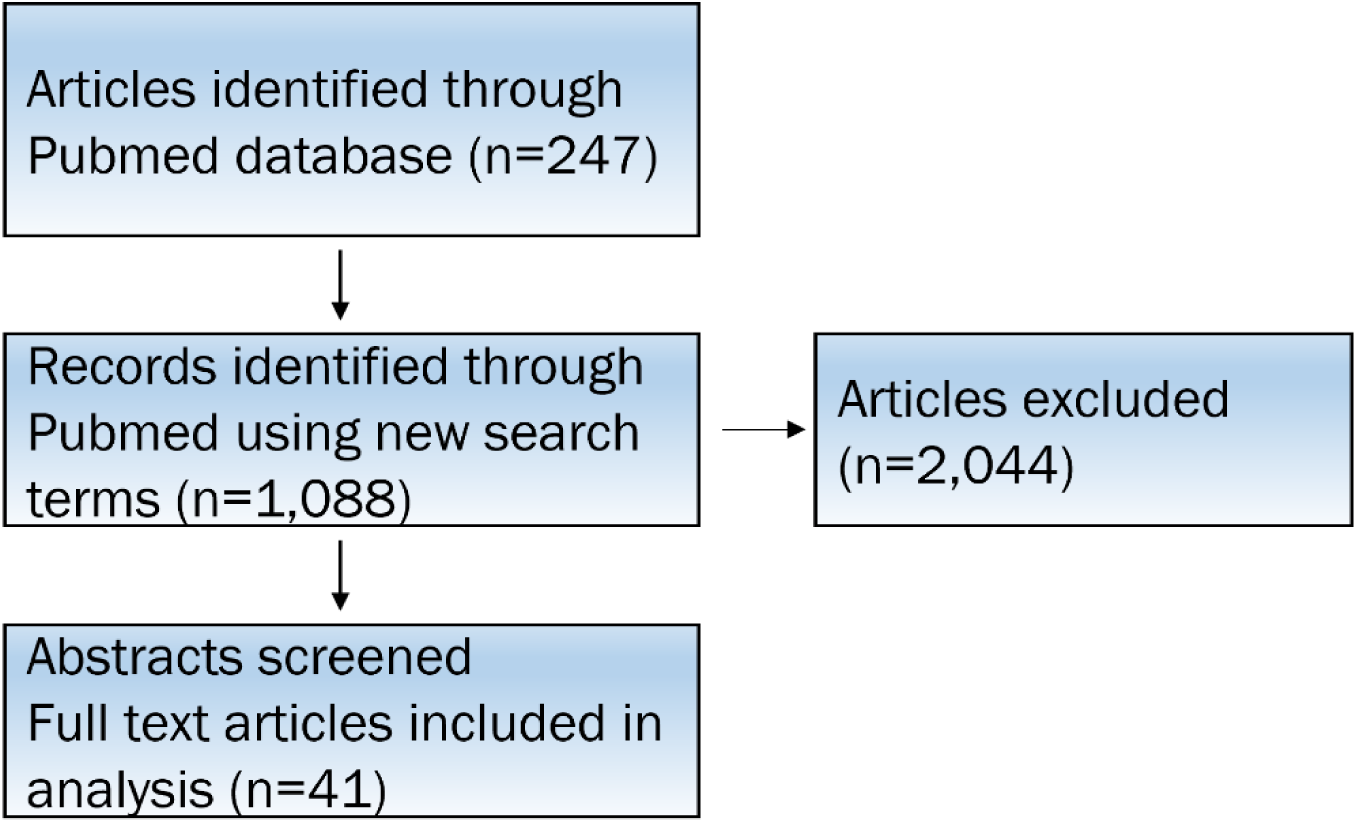
Selection of studies for inclusion in RO systematic review.

**Table S4.** Full text inclusion criteria for receptor occupancy systematic review.

| Inclusion criteria: |
| --- |
| • Drug administered systemically<br>• Reported route of administration, dose, and RO<br>• Not a review or meta-analysis<br>• English language paper<br>• Paper directly accessible from PubMed<br>• Conducted in rats, mice, or humans |

##### Data extraction and summary

From each study, data was extracted concerning the treatment, species, and recorded % receptor occupancy. Full data that were extracted are listed in **Table S5**.

**Table S5.** Data to be extracted for receptor occupancy systematic review.

| Data to be extracted: |  |
| --- | --- |
| Drug | • Drug<br>• Condition<br>• Target receptor |
| Dose |  |
| Route of administration |  |
| Species |  |
| Treatment | • Duration<br>• Frequency |
| % RO |  |

247 records from Pubmed were identified using the original search terms and 1,088 with the subsequent search terms. Of these, 41 were included in the analysis. Raw data can be found in **Data File S3**.

Of the 59 psychiatric drugs included in the search, human RO data was identified for 9 drugs (aripiprazole, clozapine, haloperidol, hydroxyzine, modafinil, olanzapine, quetiapine, risperidone, sibutramine), rat RO data for 8 drugs (amisulpride, aripiprazole, clozapine, haloperidol, methylphenidate, olanzapine, quetiapine, risperidone), and mouse RO data for 1 drug (lorazepam) (**Figure S4**).

**Figure S4.**
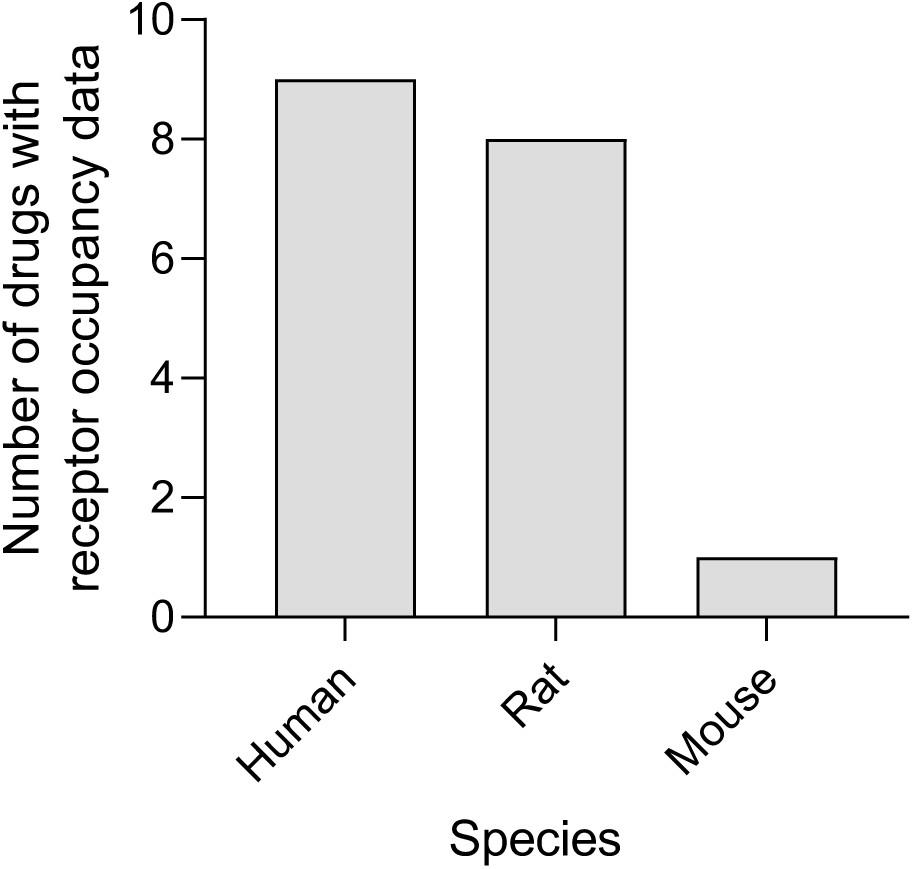
Availability of human and rodent RO data for 59 psychiatric drugs.

